# Organ structure and bacterial microbiogeography in a reproductive organ of the Hawaiian bobtail squid reveal dimensions of a defensive symbiosis

**DOI:** 10.1101/2024.11.07.622493

**Authors:** Derrick L. Kamp, Allison H. Kerwin, Sarah J. McAnulty, Spencer V. Nyholm

## Abstract

Many plants and animals house symbiotic microorganisms in specialized tissues or organs. Here, we used multidimensional *in-situ* imaging techniques to illuminate how host organ structure and bacterial microbiogeography contribute to the symbiotic function of an organ in the Hawaiian bobtail squid, *Euprymna scolopes*. Along with the well-studied light organ, female *E. scolopes* harbor a community of bacteria in the accessory nidamental gland (ANG). The ANG is a dense network of epithelium-lined tubules, some of which are dominated by a single bacterial taxon. These bacteria are deposited into squid eggs, where they defend the developing embryos from harmful biofouling. This study used a combination of imaging techniques to visualize different dimensions of the ANG and its bacterial communities. Imaging entire organs with light sheet microscopy revealed that the ANG is a composite tissue of individual, non-intersecting tubules that each harbor their own bacterial population. The organ is bisected, with tubules converging towards two points in the organ. At these points, tubules empty in a space where bacteria can mix with squid jelly to be deposited onto eggs. Observations of bacterial populations correlated bacterial taxa with cell morphology and show that tubule populations varied: some contained populations of mixed phyla while some tubules contained only one genus of bacteria. Together, these data shed light on how bacterial populations interact within the ANG and how the host uses physical structure to maintain and employ a symbiotic bacterial population in a defensive context.

**IMPORTANCE:** Sequence-based microbiome studies have revealed much about how hosts interact with communities of symbiotic microbiota, but often lack a spatial understanding of how microbes relate with each other and the host in which they reside. This study used a combination of microscopy techniques to reveal how the structure of a symbiotic organ in the female bobtail squid, *Euprymna scolopes* houses diverse, beneficial bacterial populations and deploys them for egg defense. These findings suggest that spatial partitioning may be key to harboring a diverse population of antimicrobial-producing bacteria and establish a foundation for further understanding how host structures mediate symbiotic interactions.

## INTRODUCTION

“Form follows function.” A maxim originally associated with modern architecture can be applied to a host-microbe context: physical arrangements of organs and bacterial communities affect biology and inform our understanding of how they operate. Many organisms possess symbiotic organs that have evolved to recruit and maintain microbiota with diverse functions (1–3). Structural features of these organs facilitate contact with environmental microbes during recruitment (4, 5), allow symbiont entry into the host (6, 7), or sequester symbionts to specific areas for optimal function (8–10). Disruption of this discreet localization of microbes can lead to a dysbiosis and negative impacts on host health (11–14). The microbiogeography –the spatial arrangements the bacteria have in relation to each other and host– affects the microbial partners too: spatial fragmentation of microbial populations in “microniches” can reduce competition amongst strains (15), allow specific strains to dominate an organ (16), and even drive evolution of the bacterial community (17, 18). Further, understanding the microbiogeography is vital to understanding the ecology and physiology of the entire system (19, 20).

The Hawaiian bobtail squid, *Euprymna scolopes*, is a robust model organism for studying host-microbe interactions. Primarily studied for its relationship with the bioluminescent bacterium *Vibrio fischeri*, the squid has a specialized light organ, equipped with structural features to recruit and maintain the *V. fischeri* for light production (4, 7, 21, 22). The spatial location of *V. fischeri* within the organ affects bacterial interactions (23), and the organ has physical features that optimally utilize the light produced by the bacteria (24–26). Female squid have another symbiotic organ called the accessory nidamental gland (ANG), which is part of the reproductive system. In contrast to the binary symbiosis found in the light organ, the ANG houses a diverse yet conserved community of bacteria. This consortium resides in a complex network of vascularized, epithelium-lined tubules (Fig. 1A) (27), and community analysis via 16S rRNA sequencing shows the adult ANG of *E. scolopes* is dominated by *Alphaproteobacteria* and *Verrucomicrobia*, with *Gammaproteobacteria* and *Flavobacteriia* present at lesser abundances (27–29). These bacteria are environmentally acquired by the early-stage ANG recruitment tissues that are equipped with many ciliated pores and invaginations (Fig. 1B) (28). The relative composition of this bacterial community changes as the organ develops; early-stage organs are dominated by *Verrucomicrobia* while matured organs are dominated by *Alphaproteobacteria* (30). Notably, if specific members of the bacterial community are not present in the squid’s habitat, the ANG will not form, suggesting that microbial signals trigger ANG development (31).

**Figure 1:**
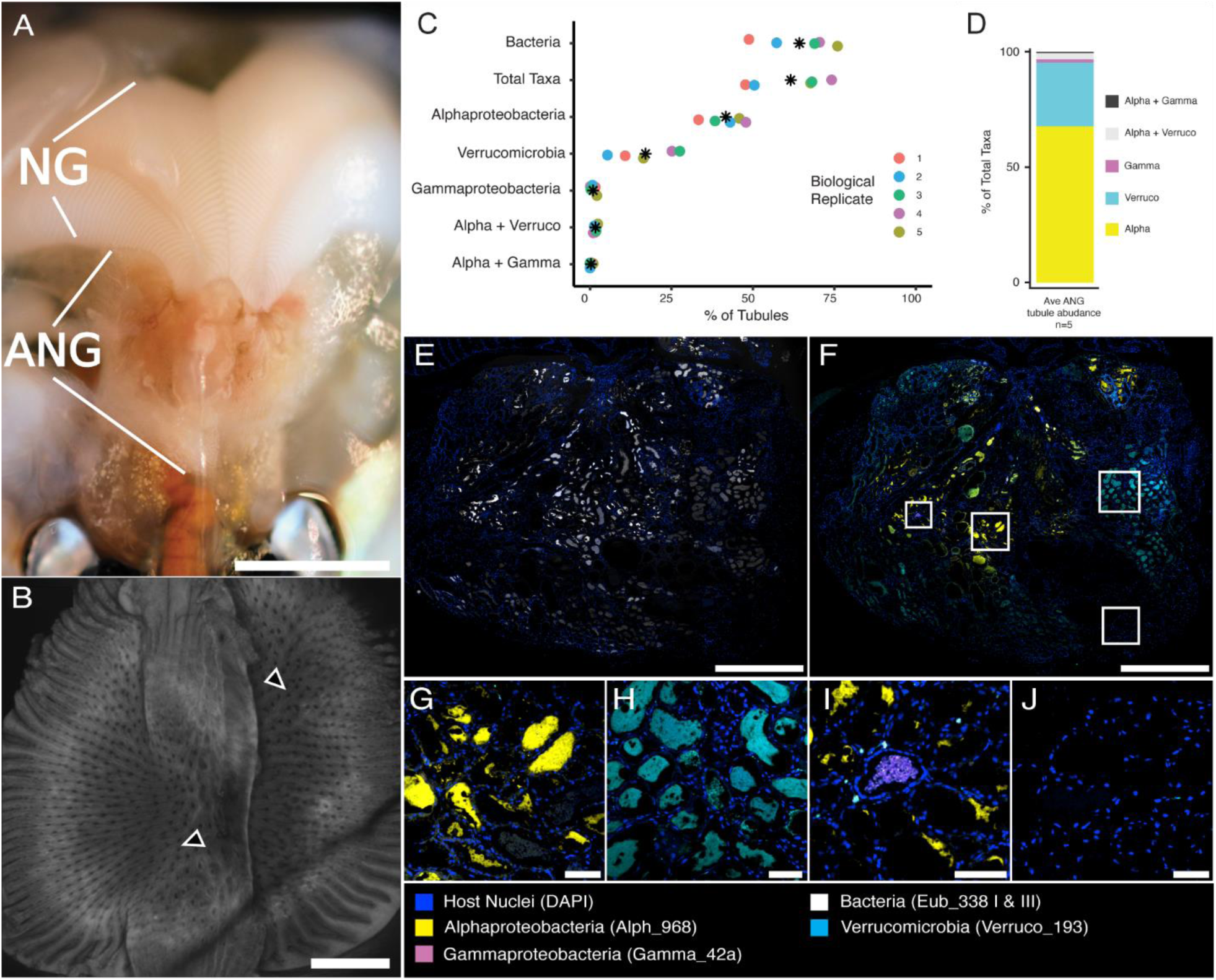
Squid anatomy and taxa abundance in the accessory nidamental gland (ANG). (A) Ventral dissection of adult *Euprymna scolopes*. The ANG lies directly adjacent to the nidamental gland (NG). (B) Early-stage ANG stained with Alexa Fluor 568-conjugated phalloidin. Recruitment tissue appears as two “pads” with many pores (black arrowheads) Adapted from (26). (C) Abundance of tubules containing each bacterial probe. “Total taxa” is the summed abundance of tubules containing any of the three represented taxa. Asterisks denote mean abundance for each group. (D) Relative abundance of tubules labeled with taxa-specific probes. (E) Fluorescence *in-situ* hybridization (FISH) image of a section of an entire ANG, labeled with near-universal bacterial FISH probe cocktail used as a positive control. (F) A section serial to panel D, showing the three most abundant higher-level bacterial taxa in the ANG. (G-J) Enlarged images of boxed regions in E, showing tubules dominated by exclusively (G) *Alphaproteobacteria*, (H) *Verrucomicrobia*, (I) *Gammaproteobacteria*, and (J) no bacteria. Scale bars: A = 1 cm; B = 200 µm E&F = 1,000 µm; G-J = 100 µm

ANGs are found in many cephalopods (32–36), and the organ has long been thought to play a role in egg laying (32, 37). The ANG lies immediately anterior to the nidamental gland (NG), an organ that produces a gelatinous jelly that coats the squid’s eggs (Fig. 1A) (34, 37). It is now understood that the ANG symbiosis is defensive: the bacteria from the ANG are transferred to the egg jelly-coat, where they produce antimicrobial compounds and protect the developing embryos from harmful biofouling by fungi and bacteria (27, 29, 38–40). Though it is known that the ANG bacteria are transferred to the egg jelly coat, the specific mechanism by which they are deposited into the jelly coat is unknown.

Sequence and culture-based techniques have revealed key elements of the ANG symbiosis. Yet the lack of spatial understanding leaves foundational questions unanswered. Previous research observed tubules that contained only one type of high-level taxa: *Alphaproteobacteria* and *Verrucomicrobia* (27). However, *Gammaproteobacteria* have not been localized in the ANG. Further, it is not understood if all tubules contain only one high-level taxon of bacteria, nor is it known how specific the taxonomic partitioning is; are bacterial populations also partitioned at lower taxonomic levels, such as genus? To date, structural understanding of ANG tissues has been limited to superficial observations or sectioned tissue, and the three-dimensional architecture of the complicated organ has remained unseen. Here, we use advanced imaging techniques to visualize the spatial organization of both a complex organ and the bacterial community that resides within. The synthesis of these data sheds light on bacterial dynamics within the ANG and informs how the host uses organ structure and spatial organization to shape a symbiotic community of bacteria involved in host defense.

## MATERIALS AND METHODS

### Animal Care and Dissection

All animal experiments were performed in compliance with protocol number A18-029 approved by the Institutional Animal Care and Use Committee, University of Connecticut. Squid were caught on shallow flats of Maunalua Bay, Ohau, HI by dip net. Animals were shipped back to The University of Connecticut and housed in aquaria for up to three months before euthanasia. Squid were anesthetized in a solution made of artificial seawater and 2-5% ethanol. Upon anesthetization, mantle lengths were recorded, animals were decapitated, and organs dissected. Whole ANGs were immediately transferred to a solution of 4% paraformaldehyde in filter sterilized squid ringers (FSSR) (27) for overnight fixation at 4°C with light rocking. Fixed tissues were washed with three consecutive washes of FSSR at room temperature and stored in FSSR at 4°C before use.

### Tissue Clearing and Whole Mount Imaging

A modified DEEP-Clear (41) technique allowed for optical clearing and depigmentation of whole ANG tissue. Briefly, fixed and washed tissues were submerged in pre-chilled acetone at -20°C for 3 hrs and washed three times with mPBS (50mM sodium phosphate buffer, 0.5M NaCl, pH 7.4) at room temperature. Samples were then treated with a solution of 75 µg/ml of proteinase K in mPBS for 12 mins at room temperature follow by two washes with a solution of 2mg/ml glycine in mPBS at room temperature. After three washes of mPBS, tissues were submerged in DEEP Solution 1.1 (10% v/v N,N,N′,N′-Tetrakis(2-hydroxyethyl)ethylenediamine, 5% v/v Triton X-100, 5% v/v Urea, UlltraPure Water, pH 10) overnight at 37°C with gentle rotation in the dark. After three mPBS washes, samples were stained with nuclear stain TOTO-3 Iodide (1 µM) and *Lens culinaris* lectin -a lectin found to universally stain tubule linings-conjugated to Rhodamine Red (20 µg/ml) in mPBS overnight with gentle rotation in the dark. After staining, samples were washed with mPBS three times at room temperature and moved back to DEEP Solution 1.1 overnight at 37°C in the dark. Samples were then washed with mPBS and submerged in a 50/50 mix of mPBS and refractive index matching media, made up of 60% w/w Opti-prep solution, 20% w/w DMSO, 20% w/w mPBS, 20 mM Tris, and 1.12 g/ml 5-(N-2-3-Dihydroxypropylacetimido)- 2,4,6-triiodo- N,N’-bis-(2,3-dihydroxypropyl) isophthalamide (Nycodenz, MP Biochemicals). After an 8 hr incubation in the dark at room temperature, samples were transferred to refractive index matching media overnight in the dark at room temperature with gentle shaking. The following day, samples were imaged in the refractive index matching media on a Zeiss Light Sheet 7. Resulting image channels were merged, false colored, and analyzed in Arivis Vision4D software (Zeiss, Germany).

### Embedding and sectioning

Fixed tissues were dehydrated in a graded ethanol series and semi-cleared with three changes of xylenes before being embedded in paraffin wax. Embedded tissues were serially sectioned into 8-10 µm thick sections on a Shandon Finesse 77510250 microtome (Thermo Scientific).

### Fluorescence *in-situ* hybridization (FISH)

Probes for targeting universal bacteria, as well as *Alphaproteobacteria*, *Gammaproteobacteria*, and *Verrucomicrobia* within the ANG were designed based on previously published literature (27, 42–44). Novel probes for targeting *Leisingera* and *Ruegeria* were designed using 16S sequence data in Arb (45) and confirmed for target specificity with TestProbe (46). Novel probes were also experimentally confirmed for target specificity and optimal formamide concentrations using three cultured representative strains of the respective genus. *Leisingera*-specific probes did not target *Ruegeria* and vice-versa. All probes were synthesized by Biomers.net (Ulm, Germany). Probe information can be found on Supplementary Table 1. Due to its limited abundance in the bacterial community and difficulty implementing into the probe set, *Flavobacteriia* were not analyzed in this study.

Sectioned tissues were deparaffinized in three changes of xylene and dehydrated in three washes of 100% ethanol. FISH probes were applied to sectioned samples at a concentration of 1 µM in a hybridization solution (900mM NaCl, 20mM Tris-HCl, 0.02% v/v sodium dodecyl sulfate, 20-30% formamide) without light in a humidifying chamber at 46°C for 3 hrs. Probes of similar formamide concentration were hybridized together. If hybridization with another formamide concentration was performed, samples were submerged in wash buffer (112-225 mM NaCl, 20mM Tris-HCl, 5mM EDTA, 0.01% v/v sodium dodecyl sulfate) at 46°C for 20 mins and the other probes and hybridization buffer were applied and washing was repeated. Slides were counterstained with 5 µg/ml DAPI (4’,6-diamidino-2-phenylindole) at room temperature for 10 mins without light. Slides were then dipped in cold Nanopure water, air dried in the dark, and mounted in Prolong Gold antifade reagent (Invitrogen) with a no. 1.5 coverslip. Mounted slides cured overnight in the dark prior to imaging.

Hybridized slides were imaged on a Nikon A1R laser scanning confocal microscope. Spectral images were acquired by sequentially illuminating the sample with 640-, 561-, 514-, 488- and 405-nm laser lines using a 40X Plan-Apochromat 1.3 NA lens or 60X Plan-Apochromat 1.4 NA lens. Linear unmixing was performed with the Nikon NIS-Elements software (Nikon) using reference spectra acquired from cultured strains hybridized with the appropriate fluorophores and imaged similarly to samples. The strongest signal for each unmixed spectra was used for each fluorophore. Unmixed images were assembled and false colored in Fiji (47). For images with dense bacterial populations, a background subtraction was performed on unmixed images to resolve individual cells. To verify spectrally acquired images, fluorophores and probe were swapped, and hybridization and imaging was repeated to recapitulate similar images (Supplemental Table 1).

To acquire larger images of whole ANG sections for quantification of tubules, we used a traditional FISH approach without spectral imaging. These slides only had FISH probes targeting *Alphaproteobacteria*, *Gammaproteobacteria*, and *Verrucomicrobia*, hybridized as described above. Eubacterial universal probes were applied to serial sections to verify bacterial presence in the taxa-specific section. Hybridized slides were imaged on a Nikon A1R laser scanning confocal microscope using a 20X Plan-Apochromat 0.75 NA lens and illuminated with 405-, 488-, 561- and 640-nm laser lines. Tubule counts were analyzed and quantified by hand in Fiji (47).

### Hematoxylin and Eosin Stain

Deparaffinized slides were rehydrated through 5 min washes in a graded ethanol series, followed by 5 mins in tap water. Sections were submerged in Gill’s 2 hematoxylin for 3 mins before gentle rinsing under tap water for 1 min. Sections were transferred to an acid alcohol differentiation solution for 1 min, rinsed again in tap water for 1 min, then stained with an Eosin Y solution for 1 min. After a 1 min rinse in tap water, sections were dehydrated again with 2 min washes in a graded ethanol series and two 3 min washes in xylene. Sections were mounted with Permount, allowed to cure overnight, and imaged on a Leica Thunder Imager with a 63X 1.40 Plan Apochromat lens. Resulting RBG images were composited and false colored in Fiji (47).

### Transmission Electron Microscopy (TEM)

Tissues were dissected, fixed, processed, and imaged by TEM exactly as previously described (27). Briefly, samples were fixed in 2.0% paraformaldehyde-2.5% glutaraldehyde in buffer A (0.1 M sodium cacodylate, 0.375 M NaCl, 1.5 mM CaCl2, 1.5 mM MgCl2, pH 7.4). Samples were washed in buffer A, then post fixed in a solution of 1% osmium tetroxide, 0.8% potassium ferricyanide, 0.1 M sodium cacodylate, 0.375 M NaCl for 1.5 h at 4°C. Upon washing and dehydration, samples were Embed 812 (Electron Microscopy Sciences) and Araldite 506 (Ernest Fulham Inc.). Thin sections were obtained using a diamond knife on a LKB Ultramicrotome V and stained with 2% uranyl acetate and Reynold’s lead citrate and viewed with an FEI Tecnai Biotwin G2 Spirit electron microscope operated at 80 kV.

## RESULTS

### Adult ANG Structure

A combination of classical and advanced light microscopy methods revealed newly described features and organization of the adult ANG tissue. Optical clearing and fluorescent staining allowed for visualization of the three-dimensional structure of the ANG tubules via light sheet microscopy (Fig. 2). In the center of the ANG, large tubules converged towards the NG (Fig. 2B). The tubules converged at two different points, each point directly underneath one half of the nidamental gland (Fig. 2E-G). In the middle of the organ, these converging tubules did not cross over the medial line of the ANG, creating the appearance of a two-lobed structure in the middle/dorsal region of the gland (Fig. 2E). Manually tracing an individual tubule from the convergence point showed that it was non-intersecting and ran continuously before it narrowed and eventually tightly bundled together in a confined space (Fig. 2A&B). As tubules reach the lateral and ventral superficial surface of the gland, the individual clustered tubules do cross the medial line and obfuscate the bilobed structure of the organ. (Fig. 2D).

**Figure 2:**
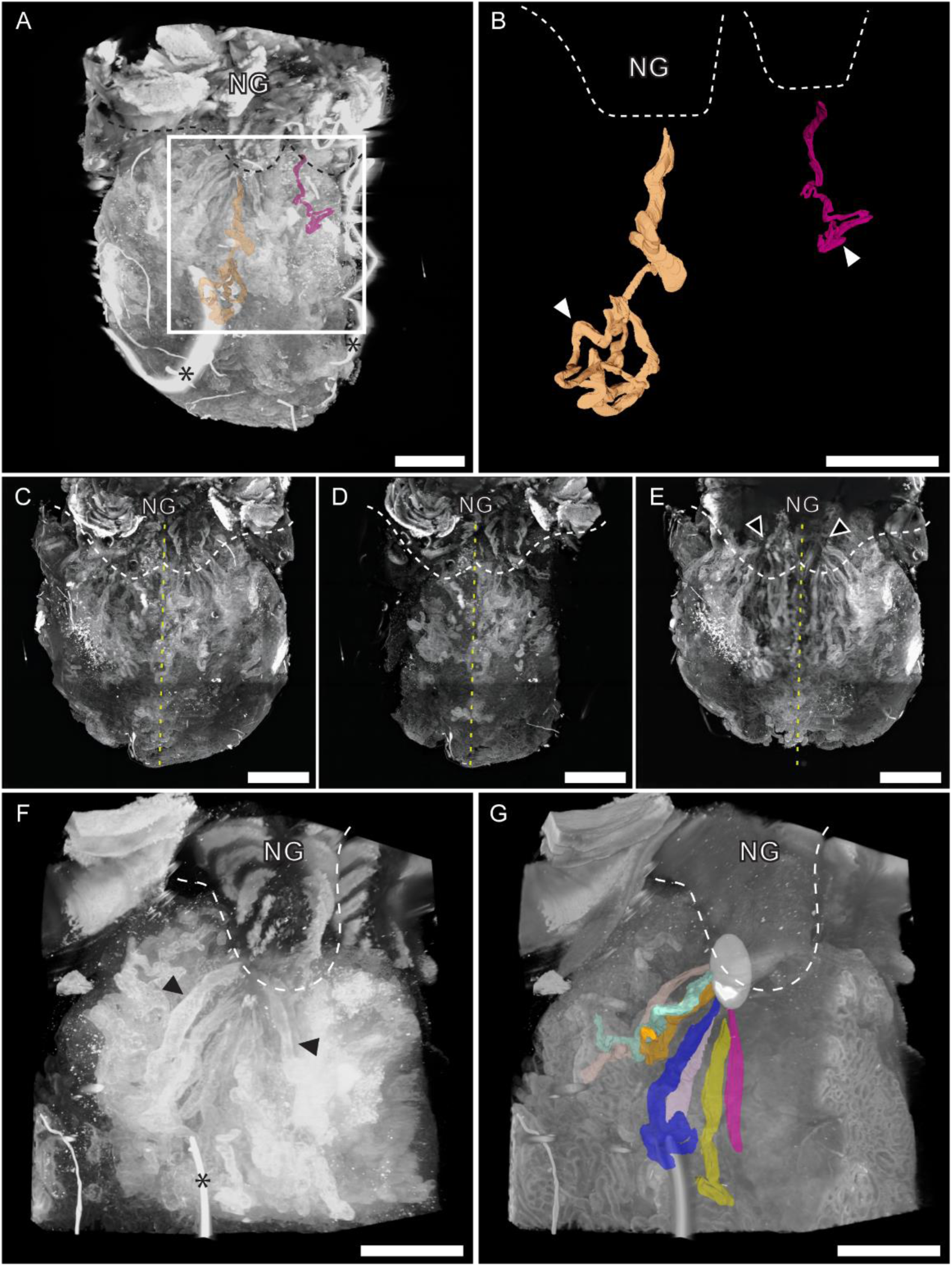
Three-dimensional organization of ANG tissue. Cleared, whole-mount tissue stained with *Lens culinaris* lectin and TOTO-3 Iodide imaged on a Zeiss Lightsheet 7 microscope. (A) Maximum intensity 3D rendering of entire ANG tissue, with representative traced tubules labeled within. (B) Enlarged image of two representative traced tubules. Individual tubules cluster in defined region, (white arrowheads), and trace continuously towards the NG. (C-E) Maximum intensity projections of the whole ANG; yellow dash represents medial line of organ. (C) The entire organ. (D) The ventral 1/3 of the organ. Superficial, ventral tubules are unorganized and cross the medial plane. (E) The dorsal 2/3 of the organ. In the middle of the ANG, tubules form a bilobed structure, and tubules converge towards a single point (black arrowheads) on each side. (F) Maximum intensity 3D rendering of a quadrant of the ANG showing individual tubules (black arrowheads). (G) Surface 3D rendering of the same quadrant in F. Representative traced tubules converge together at a point (white ovoid) dorsally adjacent to the NG. Asterisks on A and F denote dust contaminant. Scale bars: A-E = 1,000 µm; F&G = 500 µm Supplementary Videos 1 & 2

We further investigated the point where tubules converged underneath the NG. On the posterior end of the ANG, a channel in the gland creates a small space between the ANG and NG (Fig. 3A). Here, tubules terminate at pores that empty into the intergland space (Fig. 3B&C). The intergland space of one lobe was lined with approximately 125 pores, suggesting an equal number of tubules in that hemisphere of the organ. The use of FISH, we observed the three higher taxa in these converging tubules (Fig. 3D&E). Bacterial taxa observed in the ANG tubules were also observed in the intergland space (Fig. 3D-G), and bacteria-containing tubules appeared to open into the intergland space, suggesting that bacteria are secreted from the ANG into the intergland space (Fig. 3F&G). Hematoxylin and eosin (H&E) staining of tissue sections reveal that cilia line the intergland space on both the NG and ANG. (Fig. 3H&I). These cilia contained bacteria-sized particles (Fig. 3I), which were confirmed to be bacteria using FISH (Fig. 3G).

**Figure 3:**
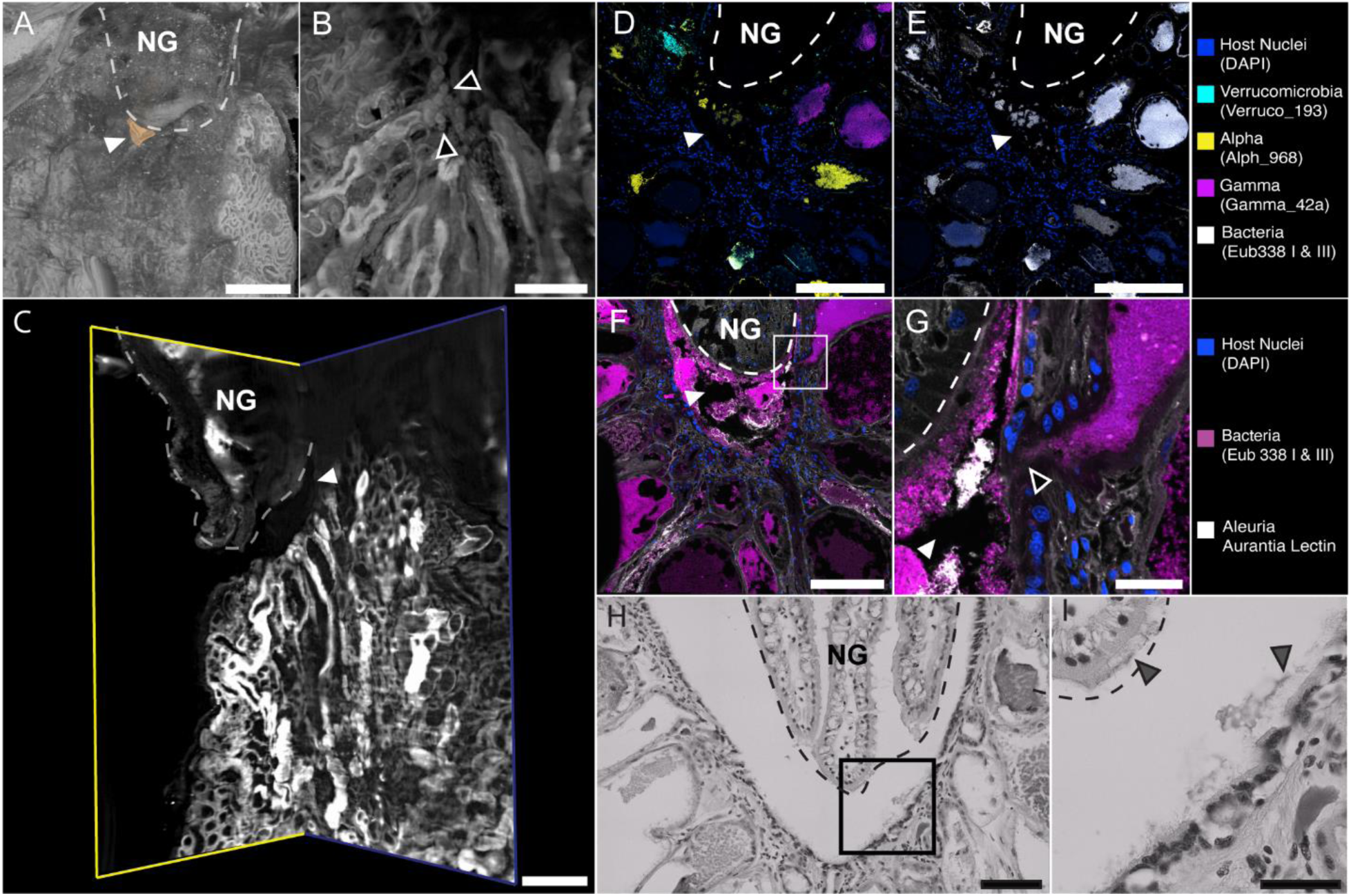
The intergland space. (A-C) Cleared, whole-mount tissue stained with *Lens culinaris lectin* and TOTO-3 Iodide imaged on a Zeiss Lightsheet 7 microscope. (A) Surface 3D rendering of ANG and NG. At the point of tubule convergence, there is an open space between the ANG and NG (white arrowhead/orange region). (B) ANG tubules terminate at pores that line the intergland space (black arrowheads). (C) Clipped surface 3D rendering of ANG and NG, showing the intergland space (white arrowhead) that opens ventrally. (D) Combinatorial labeling and spectral imaged FISH (CLASI-FISH) image of sectioned ANG tissue. The three dominant high-level taxa of the ANG are in the tubules converging at the intergland space (white arrowhead). (E) Same section as seen in panel D, labeled with near-universal bacterial FISH probe cocktail. (F) FISH image showing bacteria in converging ANG tubules and in the intergland space (white arrowhead). (G) Enlarged image of boxed region in F showing that bacteria have access to the intergland space (white arrowhead) via tubule pore (black arrowhead). (H) Hematoxylin and eosin (H&E) stained section of intergland space. (I) Enlarged image of boxed region in H. Cilia line the intergland space and contain bacteria-sized debris (grey arrowheads). Scale bars: A = 250 µm; B = 50 µm; C = 200 µm; D-F = 200 µm; G = 20 µm H = 100 µm; I = 50 µm. Supplementary Video 3

### General Distribution of Bacteria and Interactions with Host Cells

Hybridization with the near-universal Eub338I and III probe cocktail showed that a majority of tubules (64.2 ± 9.9%) contained detectable bacteria (Fig. 1). The communities of bacteria within tubule lumina were often densely populated and displayed different organizations between different tubules. Some tubules were full of bacteria, while others contained different bacterial arrangements (Fig. 1&6). These morphological differences were independent of which bacterial taxa resided within the tubule and were observed regardless of fixation method (Carnoy’s-fixed images not shown) and in whole-mount tissues, suggesting that these different arrangements were not a product of tissue sectioning.

In most samples, we observed clusters of host cells within the lumen of tubules, often toward the posterior end of the organ near the convergence point of tubules (Fig. 4). These host cell clusters contained bacteria-like debris in an eosinophilic substance, visualized on H&E-stained sections (Fig. 4A&B). The nuclei of host cells in the clusters had the hallmark horseshoe shape of hemocytes, the macrophage-like squid immune cells that circulate in the blood (48–50). FISH imaging showed that the host cells were interacting with and –in some cases– engulfing bacteria within the lumen of tubules (Fig. 4C&D). Transmission electron microscopy T also showed host cells of this hemocyte morphology interacting with bacteria inside the lumen (Fig. 4E). Imaging with taxa-specific FISH probes revealed mixtures of both *Alphaproteobacteria* and *Verrucomicrobia* were associated with these host cell clusters (Fig. 4F). *Alphaproteobacteria* and *Verrucomicrobia* were also both found in the interstitial tissue space between tubules (Fig. 4H-J). Bacteria were observed in the interstitial space throughout tissue sections. *Gammaproteobacteria* were not seen in the interstitial space.

**Figure 4:**
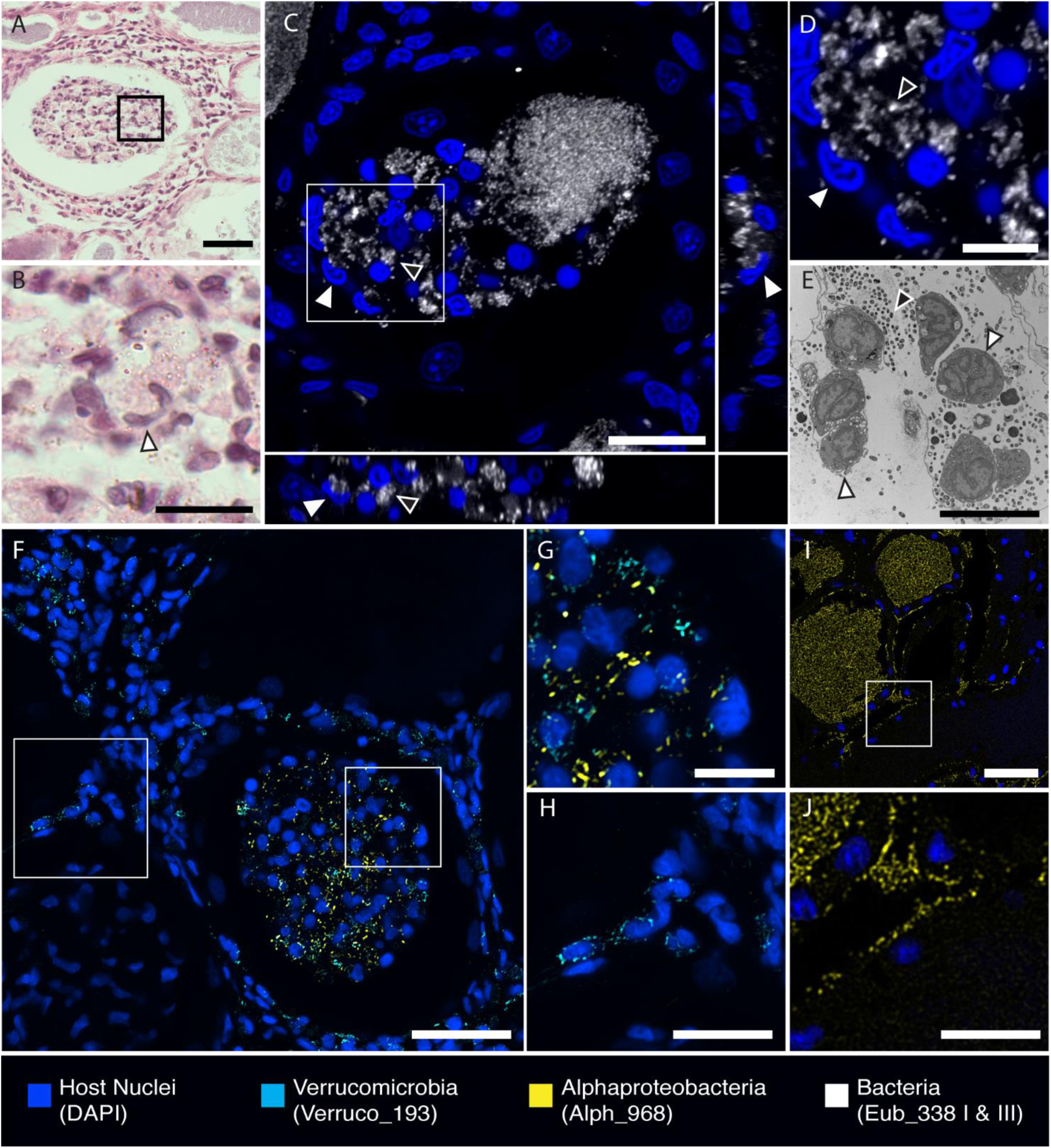
Host cells cluster in tubules and bacteria are found in interstitial tissue. (A) H&E-stained section. Host cells, bacteria, and eosin-stained material in tubules. (B) Enlarged image of boxed region in A. Host cells resemble hemocytes (white arrowheads) and surround bacteria-sized debris. (C) Orthological projection of FISH image of sectioned tissue. Hemocytes (white arrowheads) surround and engulf bacteria (black arrowheads). (D) Enlarged image of boxed region in C. (E) Transmission electron micrograph (TEM) of hemocytes (white arrowheads) inside tubules interacting with bacteria (black arrowheads). (F-G) FISH image of sectioned tissue. Bacteria in the hemocyte clusters are of mixed taxa, containing *Alphaproteobacteria* and *Verrucomicrobia*. (G) Enlarged image of boxed region in F of hemocyte cluster. (H) Enlarged image of boxed region in F, showing *Verrucomicrobia* cells in the interstitial space between tubules. (I) FISH image of sectioned tissue showing *Alphaproteobacteria* residing in tubules and in interstitial space. (J) Enlarged image of boxed region in I, showing *Alphaproteobacteria* in between tubules. Scale bars: A = 100 µm; B = 50 µm; C = 20 µm; D&E = 10 µm; F = 75 µm; G = 10 µm; H = 20 µm; I = 50 µm; J = 20 µm

### Bacterial Taxa in the ANG

We observed *Alphaproteobacteria* in all tissue sections and in the greatest number of tubules (41.7 ± 5.3%). Tubules containing *Alphaproteobacteria* were most commonly found in the center of the organ and in the tubules converging underneath the NG. These *Alphaproteobacteria* were densely packed, at times making it difficult to perceive individual cells and bacterial morphology. *Alphaproteobacteria* was the only taxa in which both cocci and bacillus bacterial morphologies were observed (Fig. 4G and 5B&C). *Alphaproteobacteria* were the most dominant members of the bacterial community secreted between the ANG and NG (Fig. 3D).

**Figure 5:**
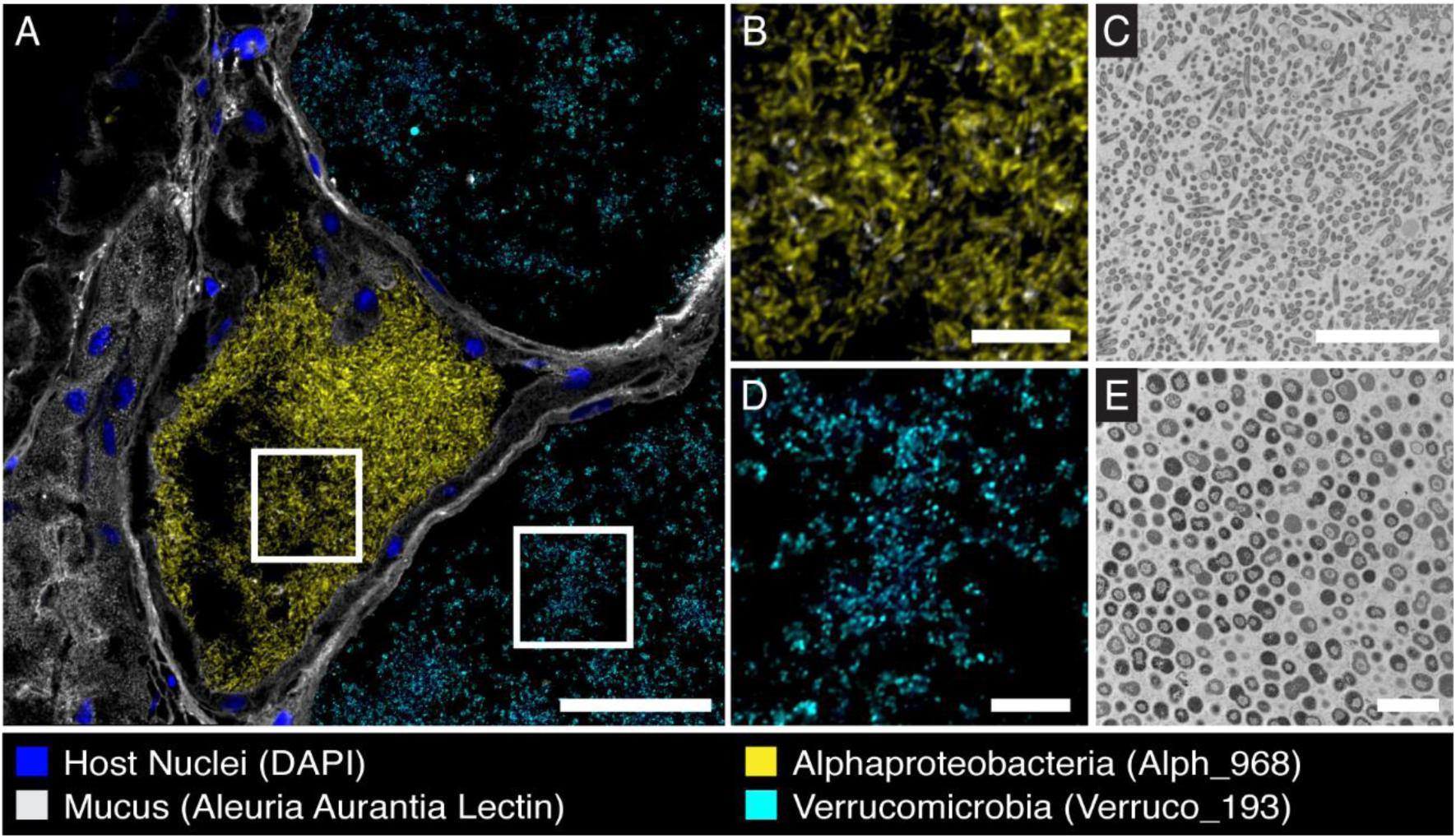
Bacterial taxa correlate with morphology. (A) FISH image of sectioned ANG tissue. (B) Enlarged image of boxed region in A. *Alphaproteobacteria* was the only taxon observed as rods. (C) TEM of rod-shaped bacteria in ANG tubules. (D) Enlarged image of boxed region in A. Verrucomicrobia were only observed as cocci. (E) TEM of cocci bacteria in ANG tubules. Scale bars: A = 75 µm; B = 15 µm; C = 10 µm; D = 15 µm; E = 5 µm

*Verrucomicrobia* were observed in the second greatest number of tubules (17.1 ± 8.4%) and found in all tissue sections. Though tubules containing *Verrucomicrobia* were located near the convergence point, they were commonly found around the periphery of the organ and contained both loosely and densely packed populations. *Verrucomicrobia* populations were only observed to be cocci morphology (Fig. 5D&E). Tubules containing *Gammaproteobacteria* were the least abundant (0.96 ± 0.77%) and were densely packed.

The CLASI-FISH imaging method allowed us to observe multiple bacterial taxa within the same tissue section (Fig. 6A-C). Though most tubules were dominated by a single taxon (Fig. 1), we also, for the first time, observed tubules with mixed populations of bacterial taxa (Fig. 6). Every organ imaged contained tubules containing mixtures of *Alphaproteobacteria* and *Gammaproteobacteria* or *Alphaproteobacteria* and *Verrucomicrobia* (Fig. 1C). These tubules were found near the center of the organ and adjacent to identically populated tubules or those dominated by exclusively *Alphaproteobacteria*. The distribution of bacteria in mixed populations varied; bacterial taxa were both homogeneously (Fig. 6D&E) and heterogeneously (Fig. 6F&G) distributed in the lumen, regardless of which taxa were present. Tubules of mixed taxa always contained *Alphaproteobacteria*. *Gammaproteobacteria* and *Verrucomicrobia* were not observed together in the same tubule. Although tubules containing mixed populations were infrequent, they were observed in every biological replicate (n=5).

**Figure 6:**
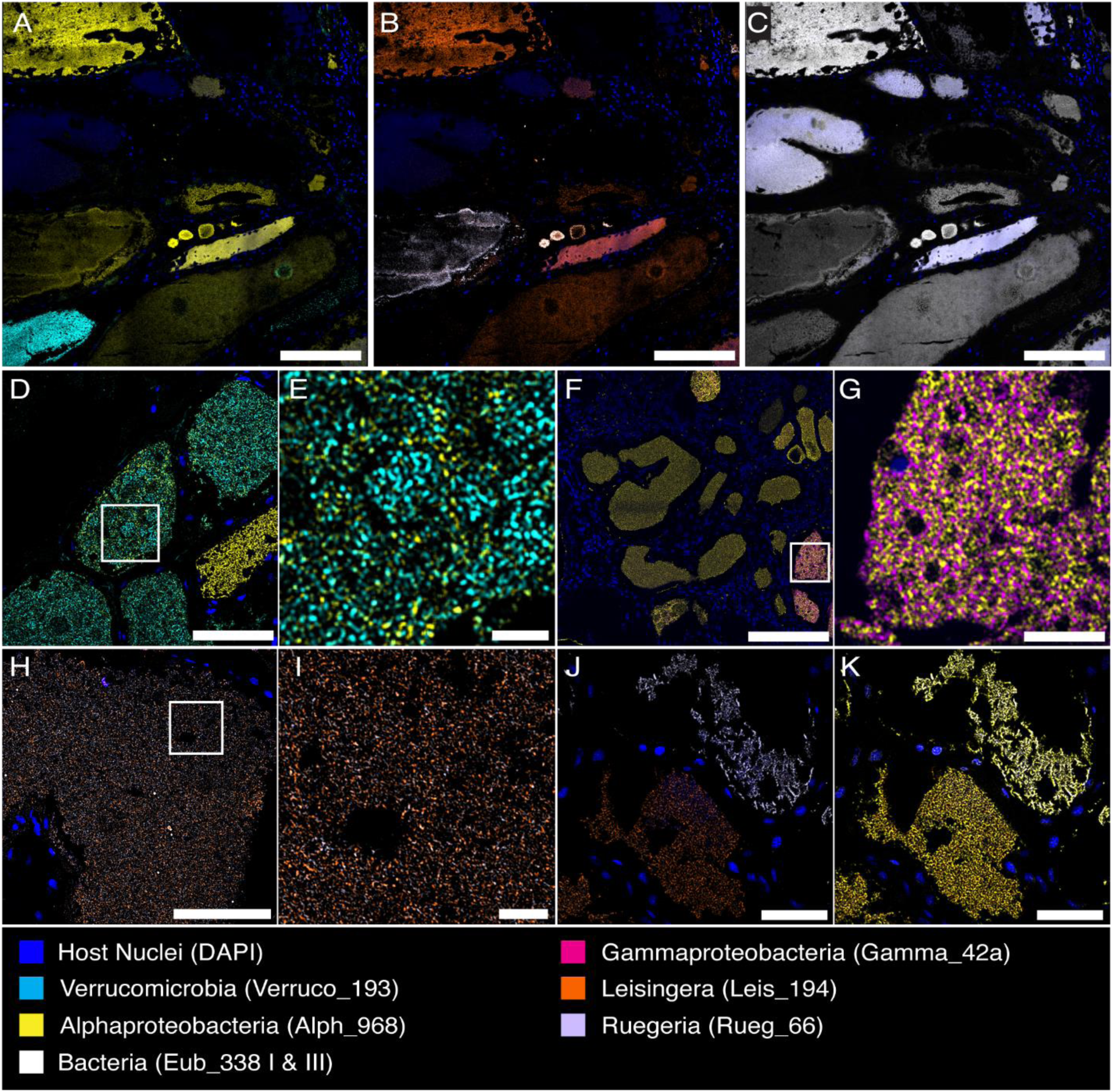
Tubules have mixed and dominant populations of bacteria at different taxonomic levels. CLASI-FISH images of sectioned tissues. (A-C) CLASI-FISH allowed for visualization of many bacterial taxa on same tissue section. (A) Higher-level taxa labeling, (B) genus-level labeling, and (C) near-universal bacteria labeling was performed on the same section. (D) *Alphaproteobacteria* and *Verrucomicrobia* in the same tubule. (E) Enlarged image of boxed region in D. (F) *Alphaproteobacteria* and *Gammaproteobacteria* in the same tubule. (G) Enlarged image of boxed region in D. (H) The genera *Leisingera* and *Ruegeria* of class *Alphaproteobacteria* were found in the same tubule. (I) Enlarged image of boxed region in H. (J) These genera were also observed as dominant in unique tubules. (K) *Alphaproteobacteria* probe overlay of J. Scale bars: A-C = 200 µm; D = 50 µm; E = 10 µm; F = 200 µm; G = 20 µm; H = 50 µm; I = 5 µm; J&K = 50 µm

Interestingly, similar observations were made with FISH probes targeting bacterial genera. In the ANG, the two most populous genera within the class *Alphaproteobacteria* are *Ruegeria* and *Leisingera* (29). These two groups were observed both individually dominating some tubules (Fig. 6J) and in mixed populations together (Fig. 6H&I). Not all tubules containing *Alphaproteobacteria* probes were labeled these genus-specific probes, suggesting that other *Alphaproteobacteria* genera reside in these tubules.

Generally, similar bacteria were seen in tubules clustered together, likely due to a tubule passing through the sectional plane multiple times as it clusters (Fig 2B). Some tubules were labeled with neither the Eub338 cocktail FISH probes nor the taxa-specific FISH probes. Tubules lacking detectable probe signal were often large, and though they were at times found at the convergence point of tubules, were most often found distal to the convergence point near the intergland space (Fig. 1J).

## DISCUSSION

Host-associated microbiome studies often focus on microbial community diversity but lack spatial context. By imaging the symbiotic bacterial community of the *E. scolopes* ANG in relation to the host, we gain a better understanding of how the host houses and uses its symbiotic microbial community. To date, structural analysis of the cephalopod ANG has been limited to two-dimensional tissue sections. By pairing these classical methods with advanced light microscopy, we report the complex structures of both host and bacterial communities in the organ.

### ANG Structure Informs Defensive Function

The complex 3D organization of tubules visualized with light sheet microscopy illuminates how bacteria are stored in the ANG and deposited onto eggs. No tubules were observed intersecting with each other, neither in whole-mount organs nor sectioned tissues. Thus, the ANG is a composite of many individual tubules, each containing its own bacterial population. The organ appears divided into two lobes, each terminating underneath the NG (Fig. 7A&E). This bi-lobed structure is superficially evident in dissections of other cephalopods; in cuttlefish there are two distinct hemispheres of the ANG (33, 35). To our knowledge, this is the first description of tubule convergence in any cephalopod ANG. The crowding of tubules on the ventral surface of the organ in *E. scolopes* previously masked this division and convergence in the adult gland. The medial division of the adult gland mirrors the bilobed, ciliated epithelial fields observed in the ANGs of *E. scolopes* earlier in ANG development (30)and suggests that the two lobes never truly merge. Rather, the tight packing of tubules forms the appearance of a single-lobed organ.

**Figure 7:**
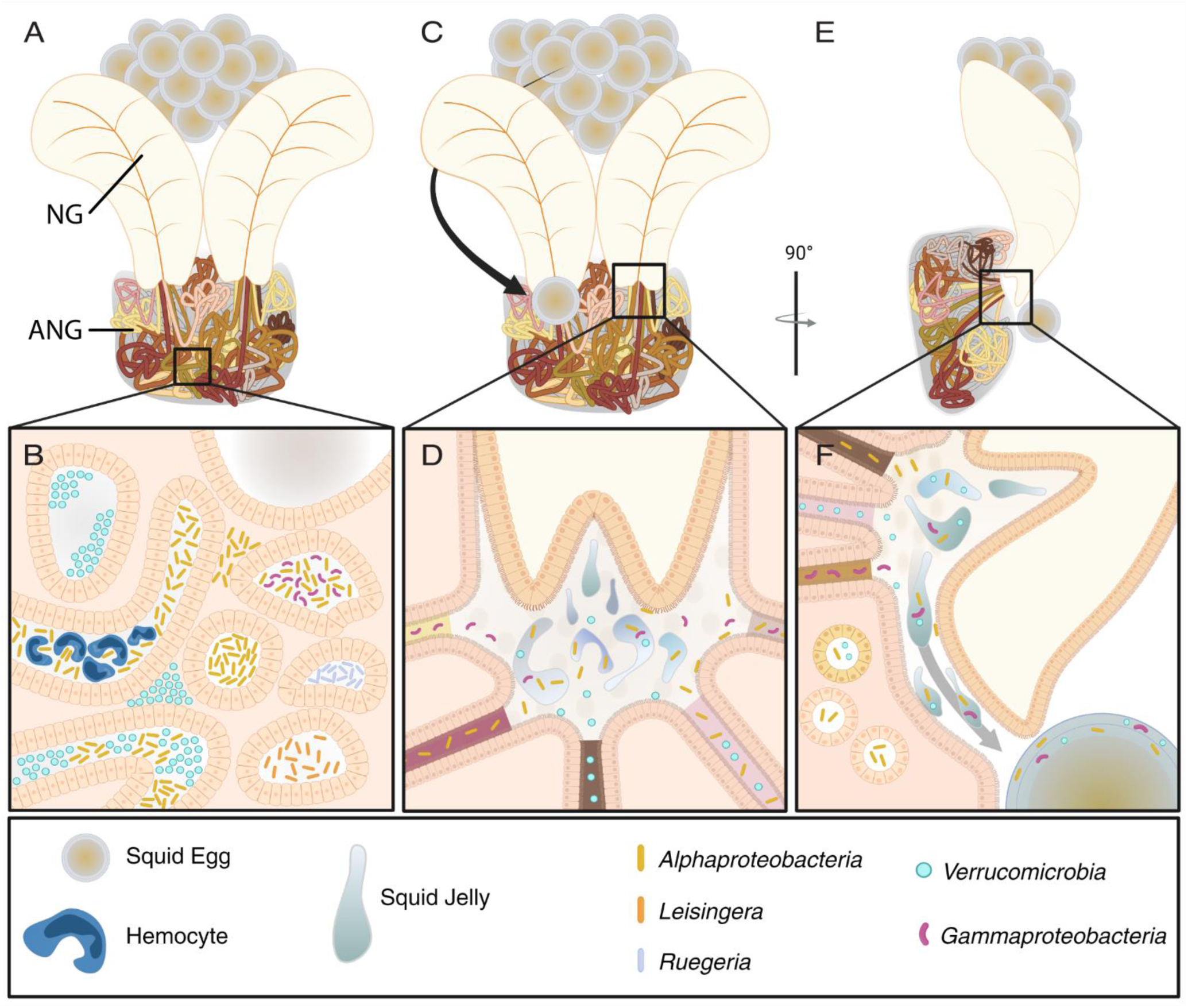
Model of ANG structure and symbiont microbiogeography during egg deposition. (A-B) Representative ANG structure and bacterial organization. (A) Individual tubules bundle together in a defined region of the organ and coverage underneath each lobe of the nidamental gland (NG). (B) The orientation of bacterial taxa and structural arrangements observed in the ANG tubules. (C-F) Proposed mechanism of bacterial deposition on eggs. (C) The egg is placed near the opening of the inter-gland space. (D) Tubules converge toward and deposit bacteria in the intergland space. Jelly from the NG is expelled into inter-gland space, and ciliary action mixes the bacteria and jelly to coat the fertilized embryo with antimicrobial-producing symbionts. (E) Rotation of organ to highlight direction of jelly and bacteria deposition during egg laying (F) Cilia transport the bacteria-jelly mixture out of the intergland space onto the squid egg.

These tubules terminate at pores lining the intergland space, which likely serves as the location where the symbiotic bacteria from the ANG mix with the jelly produced by the NG before encapsulating the egg (Fig. 7C&D). A peristaltic motion appears to push the tubule’s contents into the intergland space (Supplemental Video 4), and the cilia lining the NG and ANG in the intergland space may facilitate the mixing of bacteria and jelly as well as the transport to the egg. This mixing before egg encapsulation would allow the bacteria to be evenly deposited in the egg’s jelly coat (Fig. 7F). Other microscopic studies of ANGs observed tubules terminating and emptying at the outer surface of the ANG (34), or directly penetrating and emptying bacterial contents into the NG (35), but an intergland space has not been previously described. The three-dimensional observations of this study, together with classical tissue sectioning, suggest a mechanism by which the host squid distributes symbiotic bacteria in jelly coats of eggs during oviposition to assist in egg defense.

Symbioses involved in egg defense are prevalent in nature, and both aquatic and terrestrial animals have evolved mechanisms to employ symbiotic bacteria (51). Epibiotic bacteria associated with various crustacean eggs have been shown to protect them from fungal fouling (52–54). Though some of these studies were able to identify the specific microbes and compounds involved in the defenses (53, 54), how the host selects, maintains and deposits these symbiotic bacteria on the eggs remains unknown. Some defensive symbioses in the terrestrial environment also lack a directly observed host source for symbiont deposition: eggs of the non-biting midges *Chironomidae* contain *Firmicutes* bacteria that potentially protect the eggs from heavy metals (55), and eggs from the housefly *Musca domestica* contain bacteria that that prevent fungal fouling (56). However, organs transferring defensive symbionts have also been identified in terrestrial insects. Female beewolves, *Philanthus triangulum*, coat developing larvae with *Streptomyces* bacteria that protect the developing larva via production of antimicrobials (57–59). These symbionts are housed in specialized attennal glands and are applied to the brood pouch, where they are then transferred to larval cocoon. The beetle *Lagria villosa* has tubuled accessory glands dedicated to harboring *Burkholderia* symbionts that also protect eggs via antimicrobial production (60–62). Notably, the symbionts of these insects are vertically transmitted from mother to offspring. In contrast, the ANG microbial community is horizontally transmitted; each generation of squid acquires a new symbiotic consortium (29). Results from this study show the physical structure of the ANG facilitates the maintenance of the symbiotic microbial community, as well as the deposition of these symbionts into the eggs during oviposition.

### Bacterial Localization Expands Understanding of Host-Microbe Interactions in the ANG

The observations of the ANG bacterial community in this study support and expand upon previous understanding of symbiont localization in the ANG of *E. scolopes*. A previous study visualizing bacteria in the ANG observed both coccus and rod-shaped bacterial morphologies in tubules via TEM (27). Using FISH, we determined that both *Verrucomicrobia* and *Alphaproteobacteria* correspond with the coccus shape, but only the *Alphaproteobacteria* were rod shaped. The same study observed bacteria in the interstitial space between tubules (27), and we confirmed that both *Alphaproteobacteria* and *Verrucomicrobia* localized within the interstitial space. Both *Alphaproteobacteria* (63–66) and *Verrucomicrobia* (67–69) have been observed in intracellular spaces of other hosts ranging from protists to ticks, and future work may determine if these ANG bacteria migrate from the extracellular spaces of the tubule lumen to interstitial or even intracellular spaces. We also observed tubules with varying abundances and arrangements of bacteria, similar to what has been observed in other cephalopod ANGs (32, 34, 35, 70). The movement of bacteria from the tubules into the intergland space empties tubules and likely accounts for why some tubules appear empty at any given time. Though our study did not account for when the host last laid eggs, previous observations noted that in other squid, ANG tubules were emptied during spawning (34). Tubules that remain empty, even distal to intergland space, may have contained bacteria that have yet to repopulate the tubule, such as the putatively slow growing *Verrucomicrobia* (71).

The relative abundance of tubules containing each taxon of bacteria resembled the relative abundance of each taxon as estimated via 16S community analysis (29), suggesting that spatial structuring governs bacterial abundance in the organ: the more tubules containing a given bacterial taxon, the more that taxon will be represented in the ANG community. The ANG houses a diverse community of bacteria that produces a broad array of antimicrobial products that protect the squid eggs from fouling pathogens (3, 38, 39). Previous work showed the complete partitioning of bacterial taxa into separate tubules (27), and similar observations were made in this study (Fig. 1 G-J). This spatial partitioning may maintain the diversity of antimicrobial-producing bacteria. Spatial segregation of bacteria increases the biodiversity in mock experimental communities dominated by negative interactions (72), and host-driven physical structuring of microbial communities on the human skin allows dissimilar strains of bacteria to coexist in the same organ by reducing competition (15). This competition reduction may most benefit microbial taxa that are rare in the host’s environment, such as the *Verrucomicrobia* which are of low relative abundance in the bobtail squid’s habitat (0.3%) but are one of the dominant taxa in the ANG (25%) (29). Many defensive symbioses involve a limited diversity of bacterial members (51, 57, 61, 62). But in the ANG, spatial partitioning lessens both negative interactions and competition between bacterial community members, thus allowing for a diverse consortium of antimicrobial-producing bacteria.

A variety of mechanisms may work in concert to maintain a spatially partitioned bacterial population. The conserved bacterial communities between individual host ANGs suggests that the host actively selects for specific bacteria from the environment. We observed macrophage-like hemocytes interacting with bacteria in ANG tubules, similar to what has been observed in a previous study (27). Hemocytes are also known to migrate to the light organ, where they interact with *V. fischeri* and play multiple roles in the light organ symbiosis (49, 73–76). In the ANG, they may perform similar functions, including restructuring tissues, regulating symbionts via binding and phagocytosis, or delivering nutrients to the symbionts. Notably, the hemocyte aggregations with eosinophilic material surrounding bacteria in the ANG resemble hemocytes that respond to parasites and pathogens in other invertebrates (77–79), suggesting that the host’s cellular innate immune system may help regulate the consortium, possibly by removing unwanted bacteria from tubules. Further, individual tubules may foster microenvironments that lend competitive advantage to specific target bacterial taxa, similar to how different bean bugs select for their respective *Burkholderia* symbionts (80). Bacterial interactions may also lead specific taxa to dominate an individual tubule. Bacteria in the *E. scolopes* ANG produce antimicrobial compounds (38, 39), and extracts from other cephalopod ANGs have demonstrated antibacterial activity (70, 81). Further testing may reveal if bacteria producing these antimicrobial compounds dominate individual tubules by eliminating other members via antimicrobial production. Many ANG bacteria also possess type VI secretion systems (82) and may use them to eliminate bacterial competitors in tubules, just as some *V. fischeri* strains do in the light organ crypts (83). These potential inhibitory interactions emphasize the need for the host to partition the bacterial community. Experiments focused on coculturing ANG bacteria with other bacterial strains or the isolatable squid hemocytes (50) will potentially reveal the mechanisms underlying these interactions. The use of more FISH probes on entire tissue sections revealed that some tubules contain more than one bacterial taxon (Fig. 6). The bacterial interactions in these mixed-population tubules could be syntrophic, such as those observed with mucus-degrading bacteria in the human and insect guts (84–86). However, these inter-bacterial relations may also be antagonistic. The microbial community of the ANG is originally dominated by *Verrucomicrobia* early in development but is later dominated by *Alphaproteobacteria* (30). These mixed tubules may result from direct interaction between community members, e.g. *Alphaproteobacteria* invading tubules of other bacteria, perhaps by migrating through the interstitial region between tubules or from the intergland space. This spatial localization has physiological implications: the high density at which *Alphaproteobacteria* bacteria inhabit tubules, along with their ability to cohabit with or invade tubules of different bacteria, likely accounts for the group’s predominance in the adult ANG bacterial community.

Understanding the paradox of how some tubules contain individual genera while others contain mixed phyla may also require further study of the early-stage recruitment tissues. The nascent ANG has many small pores primed to recruit environmental bacteria, but the mechanisms by which the host selects and partitions its specific bacterial consortium are currently unknown. One hypothesis is that stochastic mechanisms, such as ecological drift and the bottlenecking of bacterial populations, drive the partitioning of bacterial consortia. In human skin, small pores fragment populations of *Cutibacterium* via neutral bottlenecking (15) and in the squash bug, *Anasa tristis*, symbionts are randomly compartmentalized into different symbiotic crypts (87). Partitioning in the ANG may be driven by similar stochastic effects during colonization. The approximately 700 small pores of the ANG recruitment tissue may be individual bottlenecks that permit very few bacterial cells to enter any given pore (30). Competition between different bacterial groups and/or host factors such as innate immunity or different nutrient microenvironments may then select for the expansion of these primary bacterial symbionts and maintain partitioning of the community in developing tubules. In some cases, these primary symbionts may be ANG-compatible bacteria but belong to different taxonomic groups. These ANG-compatible bacterial populations may be bacteria from the same taxonomic class, as observed with the *Ruegeria* and *Leisingera* mixed tubules, or from different taxonomic phyla, as seen with the *Alphaproteobacteria* and *Verrucomicrobia* mixed tubules. The combination of the host-derived selection and bacteria-bacteria inhibitory interactions may result in the relative infrequency of tubules observed with mixed bacterial populations. Experiments in both the bobtail squid light organ and ANG show that the host responds to the presence of specific environmental bacteria (7, 31, 76, 88). Thus, it may be that the initial bacterial community is partitioned in the recruitment tissue pores, but the host’s selective maintenance is tailored in response to the specific bacterial population within the developing tubule. Further imaging of the nascent ANG will reveal if and how these spatial divisions structure the adult ANG symbiotic community.

By visualizing organ structure and microbiogeography of bacterial symbionts in the *E. scolopes* ANG, this study revealed how the host fosters and regulates its symbiont community. Three-dimensional imaging clarified the organization of the ANG tubules and revealed direct mechanisms used by the host to deposit the bacteria into the eggs where the bacteria provide antimicrobial protection. In all, the bobtail squid ANG represents a composite system consisting of individual tubules harboring distinct populations of bacteria, some of which are dominated by one bacterial taxon while a few contain a mixture of different taxa. We propose that partitioning the bacterial population allows the host to foster a greater diversity of antimicrobial-producing bacteria. Using multidimensional imaging methods, we show that bacterial populations are localized and mappable in a symbiotic organ, and future research correlating bacterial taxa with respective host microenvironments may reveal the cellular and biochemical mechanisms used to select and maintain specific bacteria. The foundation of this study, along with the ability to experimentally introduce bacteria into the developing organ (31), supports the ANG symbiosis in *E. scolopes* as a robust model for understanding the regulation of bacterial consortia in defensive symbioses.

## Acknowledgments

This research was conducted on the traditional and unceded lands of the Mohegan, Pequot, Nipmuc, Wôpanâak and Kānaka Maoli peoples. We honor elders past and present.

We thank Jessica Mark-Welch for assistance with CLASI-FISH and Christopher O’Connell and Emery Ng for microscopy assistance. Andrew Collins assisted with EM images and gave helpful manuscript comments. We thank Sarah Provencher, Sila Inanoglu, Esme Ho, Aaron Richardson, McKenna Oberheim, Kira O’Brien, Niko DeSousa, and Marylin Salvador for their indelible help with animal husbandry. This work was supported by NSF-IOS 2247195 and the Gordon and Betty Moore Foundation to SVN. We have no conflicts of interest to declare.

